# Characterizing and Mitigating Protocol-Dependent Gene Expression Bias in 3′ and 5′ Single-Cell RNA Sequencing

**DOI:** 10.64898/2026.02.28.708722

**Authors:** Valeryia Shydlouskaya, S. M. Mansour Haeryfar, Tallulah S. Andrews

## Abstract

Single-cell RNA sequencing (scRNA-seq) has enabled large-scale characterization of cellular heterogeneity; yet, integrating datasets generated through different library preparation protocols remains challenging. For instance, comparisons between 10X Genomics 3′ and 5′ chemistries are complicated by protocol-dependent technical biases imposed by differences in transcript end capture and amplification. While normalization, and often batch correction, is an integral step in preprocessing scRNA-seq datasets, it remains unclear which correction is most appropriate, or even necessary, for reliable cross-protocol comparisons. Here, we systematically characterize protocol-related expression differences using 35 matched donors across six tissues profiled with both 3′ and 5′ scRNA-seq approaches. We find that gene expression discrepancies are not pervasive across the whole transcriptome, but driven instead by a relatively small, reproducible subset of protocol-biased genes. Excluding these genes improves cross-protocol concordance, indicating that most genes are directly comparable without aggressive correction. We then benchmark commonly employed normalization approaches and show that while several methods, such as fastMNN, improve statistical alignment when cell populations are well matched, they can distort gene-level signals and inflate differential expression in biologically realistic settings with incomplete cell-type overlap. Taken together, our results demonstrate that protocol bias between 3′ and 5′ scRNA-seq is limited in scope and that targeted handling of a small set of biased genes presents an alternative approach to normalization or batch correction strategies. This work provides a practical guideline for integrating 3′ and 5′ scRNA-seq data and highlights the importance of matching normalization strategies to the structure of technical variation and the intended downstream analyses.

## Introduction

Single-cell RNA sequencing (scRNA-seq) is a recent technology that has become an invaluable tool for gene expression analysis with single-cell resolution. In recent years, this technique has been widely used to address key biological questions of cell heterogeneity in different tissues and disease states, including but not limited to cell differentiation, immune responses, and tumor heterogeneity [1–6]. Current scRNA-seq technologies require single cell isolation, capturing or tagging of mRNA with unique barcodes, and cDNA synthesis, fragmentation and amplification to generate sequencing libraries consisting of ∼300-base-pair-long barcoded transcript fragments. Due to the necessity of adding unique molecular identifiers (UMIs) during the reverse transcription step to reduce amplification noise [7,8], barcodes are attached to only one end of each transcript, either 3′ or 5′, which means only the fragments containing the appropriate end are amplified and sequenced.

Originally, scRNA-seq protocols included barcodes on the poly-T oligo nucleotides used to capture the mRNA molecules. As a result, UMIs were attached to the 3′ end of each transcript during reverse transcription [9,10]. However, this approach was far from ideal in interrogating immune landscapes shaped by T and B lymphocytes since only the constant region of recombined T-cell receptors (TCRs) and B-cell receptors (BCRs) could be sequenced. To enable single-cell resolution characterization of the recombined and variable portions of TCRs and BCRs, protocols were altered to instead include UMIs and cell-specific barcodes in switch oligonucleotides, thereby adding them to the transcripts’ 5′ end [11]. Accordingly, many newer scRNA-seq studies, particularly those focusing on immune landscapes and responses, use this 5′ scRNA-seq protocol [12–14]. It is noteworthy, however, that this approach has complicated the efforts to generate comprehensive atlases or perform meta-analyses due to technical biases in gene expression measured with 3′ and 5′ technologies [15].

Previous benchmarking studies have examined the efficacy of different computational methods at integrating various protocols in a shared lower-dimensional space [15–17]. However, these approaches are limited to improving the results of clustering, trajectory inference, or other algorithms that only utilize a lower dimensional representation of scRNA-seq data. They do not enable comparisons of gene expression levels between different cell populations or biological conditions, including between healthy and diseased states. To date, no systematic comparisons have been conducted to evaluate available approaches for gene-expression space normalization to correct technical differences across scRNA-seq protocols. Moreover, protocol-related differences between scRNA-seq and single-nucleus RNA sequencing (snRNA-seq) have been assessed, demonstrating that certain technical effects can be partially corrected through simple adjustments like regressing out gene length [18,19]. However, this approach does not fully reconcile scRNA-seq and snRNA-seq data due to confounding biological effects, such as cytoplasmic post-transcriptional regulation [20] and dissociation-associated stress responses that are more common in scRNA-seq than in snRNA-seq [21]. Differences between 10X Genomics 3′ and 5′ scRNA-seq protocols are simpler and exclusively result from technical variations in transcript end capture during reverse transcription and downstream amplification [22]. As these protocol-related effects are not expected to reflect genuine biological differences between cells, they may be well suited to correction using existing normalization and batch correction approaches. Nonetheless, it remains unclear which methodologies, if at all needed, are most effective at correcting 3′ and 5′ scRNA-seq data without distorting biologically meaningful signals.

To address the above gap, we selected a set of widely used normalization and batch correction methods that explicitly operate at the level of gene expression values and span diverse algorithmic approaches [15,17]. These include linear adjustment methods, such as ComBat [23] and limma [24]; mutual nearest neighbor–based approaches, including mnnCorrect [25] and fastMNN [25], which adjust expression values by aligning shared cellular neighborhoods across batches; and model-based, gene-wise normalization frameworks such as M3Drop [26] and SCTransform [27], which use negative binomial models to account for depth-dependent technical noise and produce Pearson residuals that can be used as corrected expression values. We also evaluated a simple rescaling approach based on Z-transformation, as well as deep generative models such as scVI [28] and scArches [29], which learn latent representations of scRNA-seq data and which can reconstruct normalized expression estimates while accounting for technical variation.

We have now characterized protocol-dependent gene expression differences between 3′ and 5′ scRNA-seq and evaluated the performance of existing batch correction methods to remove such differences. Using data from 35 subjects from six publicly available datasets with matched 3′ and 5′ scRNA-seq, we first identified a subset of genes whose expression is consistently biased between these chemistries. We then evaluated 10 selected correction methods for their ability to mitigate the noted biases. Performance was assessed by comparing the similarity of cell-type-specific expression profiles between 3′ and 5′ data, evaluating the degree of mixing between 3′ and 5′ samples following correction, and examining the agreement in cell-type-specific marker genes. Finally, we test the top-performing methods in a synthetic use-case designed to integrate and compare partially overlapping 3′ and 5′ datasets. Based on these results, we provide actionable guidelines on when correction is necessary and which approaches are most appropriate for integrating 3′ and 5′ scRNA-seq data in biologically meaningful ways.

## Methods

### Dataset acquisition

Six human datasets were identified from the CellxGene database that contained the same biological samples profiled using both 5′ and 3′ Chromium scRNA-seq (Table 1) [30–36]. All processed datasets and their matching metadata, including cell-type annotations, were downloaded. For each dataset, we identified subjects with matched 3′ and 5′ scRNA-seq samples generated from the same tissue using the same cell isolation strategy, when applicable. To ensure consistent quality between 5′ and 3′ data, any donors with >25% discrepancy in their frequency of any cell types between 3′ and 5′ samples, where the cell type in question represented at least 5% of the total dataset, were excluded (Fig. S1). This was with the exception of the datasets generated by Andrews [34] and Suo (thymus) [33], which were intentionally selected to represent more biologically imbalanced and, therefore, challenging scenarios. To maximize clarity and reproducibility, we mostly used the cell-type annotations provided by the dataset authors, recognizing that alternative immunological classifications may be possible. Datasets with fine-grain annotations that identified more than 20 cell types, many of which were highly similar, were manually re-annotated at a coarser level, accounting for cell-type frequencies [32–34,36] (Tables S1-S4). Cells from rare cell types (<50 cells in either 3′ or 5′ samples) were removed to prevent sparsely represented populations from biasing normalization and integration performance assessments.

**Table 1.** Description of datasets used in this study.

| Dataset | Description | Tissue(s) | Technologies/<br>Batches | Subjects Chosen | Total<br>Cells | CellXGene<br>accession link |
| --- | --- | --- | --- | --- | --- | --- |
| Park [31] | Human Thymic Development | Thymus | 10x 3' v2,<br>10x 5' v1 | "F29 45P", "F30 45P",<br>"F38 45P", "F41 45P",<br>"F41 45N", "F45 45P",<br>"F45 45N", "P1 CD3P",<br>"P1 CD3N" | 39,709 | https://cellxgene.cziscience.com/collections/de13e3e2-23b6-40ed-a413-e9e12d7d3910 |
| Suo [33] | Human Immune System Across Organs | Bone marrow | 10x 3' v2,<br>10x 5' v1 | "F41", "F45" | 18,793 | https://cellxgene.cziscience.com/collections/b1a879f6-5638-48d3-8f64-f6592c1b1561 |
|  |  | Kidney |  | "F41" | 6,006 |  |
|  |  | Liver |  | "F32", "F34", "F38",<br>"F41", "F45" | 25,983 |  |
|  |  | Spleen |  | "F45" | 17,187 |  |
|  |  | Thymus |  | "F38", "F41", "F45" | 32,673 |  |
| Andrews [34] | Human Liver | Liver | 10x 3' v2,<br>10x 5' v1 | "C59", "C61",<br>"C63", "C64" | 22,354 | https://cellxgene.cziscience.com/collections/0c8a364b-97b5-4cc8-a593-23c38c6f0ac5 |
| Simone [35] | Human Peripheral Blood Mononuclear Cells (PBMCs) | PBMCs | 10x 3' v3,<br>10x 5' v2 | 4 samples (2 repeats per batch) from 1 subject | 29,944 | https://cellxgene.cziscience.com/collections/398e34a9-8736-4b27-a9a7-31a47a67f446 |
| Leader [36] | Human Non-Small Cell Lung Cancer | Lung (malignant and normal adjacent) | 10x 3' v2,<br>10x 5' v1 | 1 donor with two tissue samples:<br>"Leader_Merad_2021_522_normal_adjacent", | 11,755 | https://cellxgene.cziscience.com/collections/edb893ee-4066-4128-9aec- |
|  |  |  |  | “Leader_Merad_2021_522_tumor_primary” |  | 5eb2b03f8287 |
| Jardine [32] | Fetal Bone Marrow | Bone marrow | 10x 3' v2,<br>10x 5' v1 | “F30”, “F41”, “F45” | 11,569 | https://cellxgene.cziscience.com/collections/793fdaec-5067-428a-a9db-ecfe135c945 |

### Data preprocessing and highly variable genes

Datasets were split into pairs of matching 3′ and 5′ data for each subject and processed separately, preventing genetic differences between individuals from confounding the results. Raw counts were log2-cpm normalized using Seurat (v5.1.0) [37] with a 10,000 counts/cell default and a pseudocount of 1. The top 3,000 highly variable genes (HVGs) were identified using a vst model [27]. All samples originating from the same tissue within each dataset were combined, and a consensus set of 3,000 HVGs was identified using Seurat.

### Identification of protocol-biased genes

To identify genes that were consistently differentially expressed between the 10X 3′ and 10X 5′ assays, we conducted differential gene expression analysis for each subject across all cells using a Wilcoxon rank-sum test implemented in Seurat (v5.1.0). We then calculated the cumulative number of overlapping differentially expressed genes (DEGs) across subjects at three different *p*-value thresholds (*p* < 1e^-5^, 1e^-50^, and 1e^-100^). Consensus biased genes were identified at the “elbow” in the cumulative distribution. Uniform Manifold Approximation and Projections (UMAPs) were generated using default parameters with 30 principal components before and after the removal of consensus biased genes. They were further inspected for integration effectiveness using Seurat ‘MixingMetric’ [38] with a neighbourhood size of 150 cells to balance between looking at local and global scales. Consequently, a consensus list of 867 protocol-biased genes (significant in at least 5 donors at 1e^-50^) was selected for downstream analyses. To assess the contribution of these genes to platform differences, we computed cosine similarity scores between 3′ and 5′ data using the biased gene list and three replicates of randomly sampled gene sets of equal size and expression level composition.

### Data normalization

Each individual pair of 3′ and 5′ data was normalized using the following approaches (Table 2):

**Table 2.**
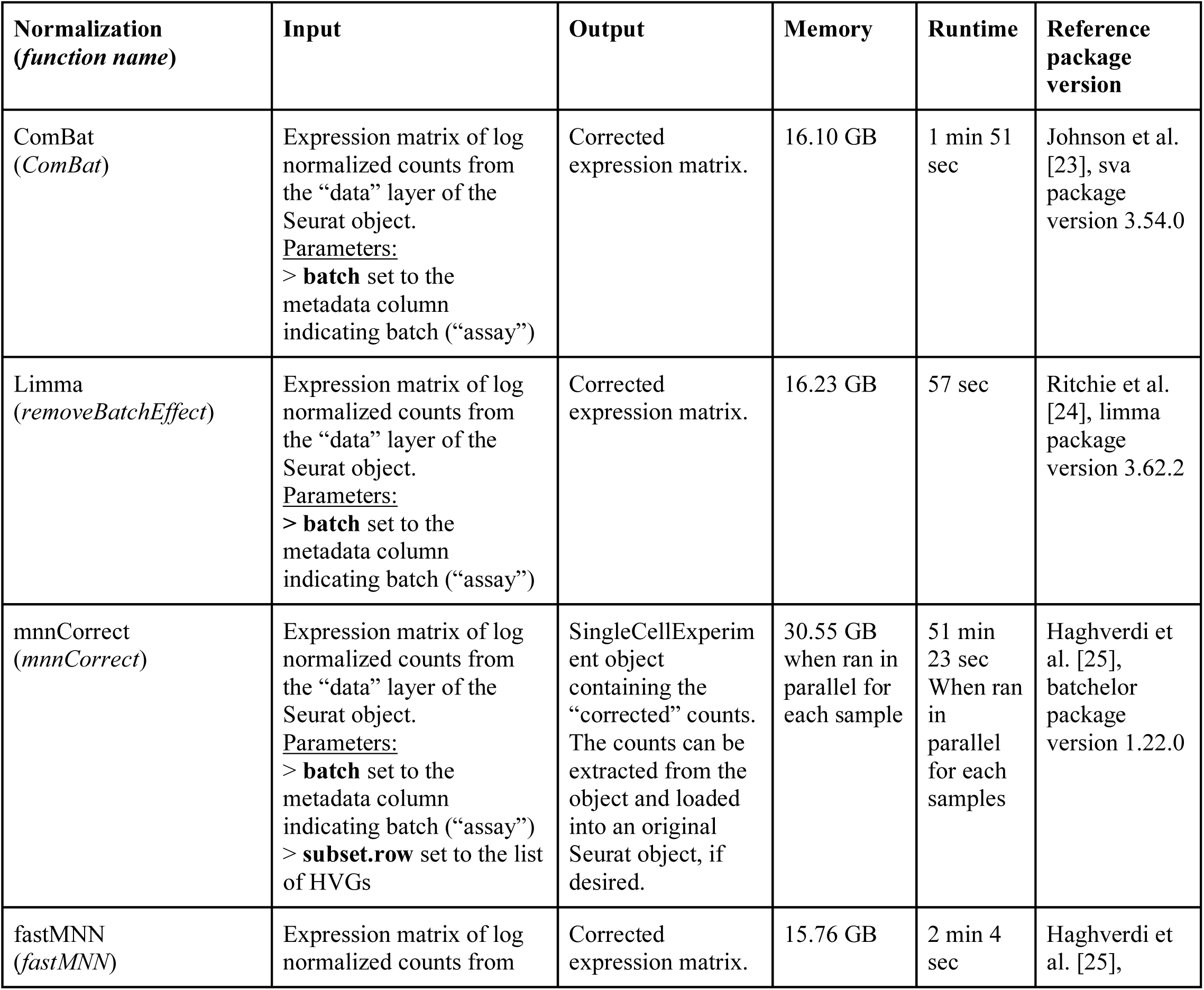
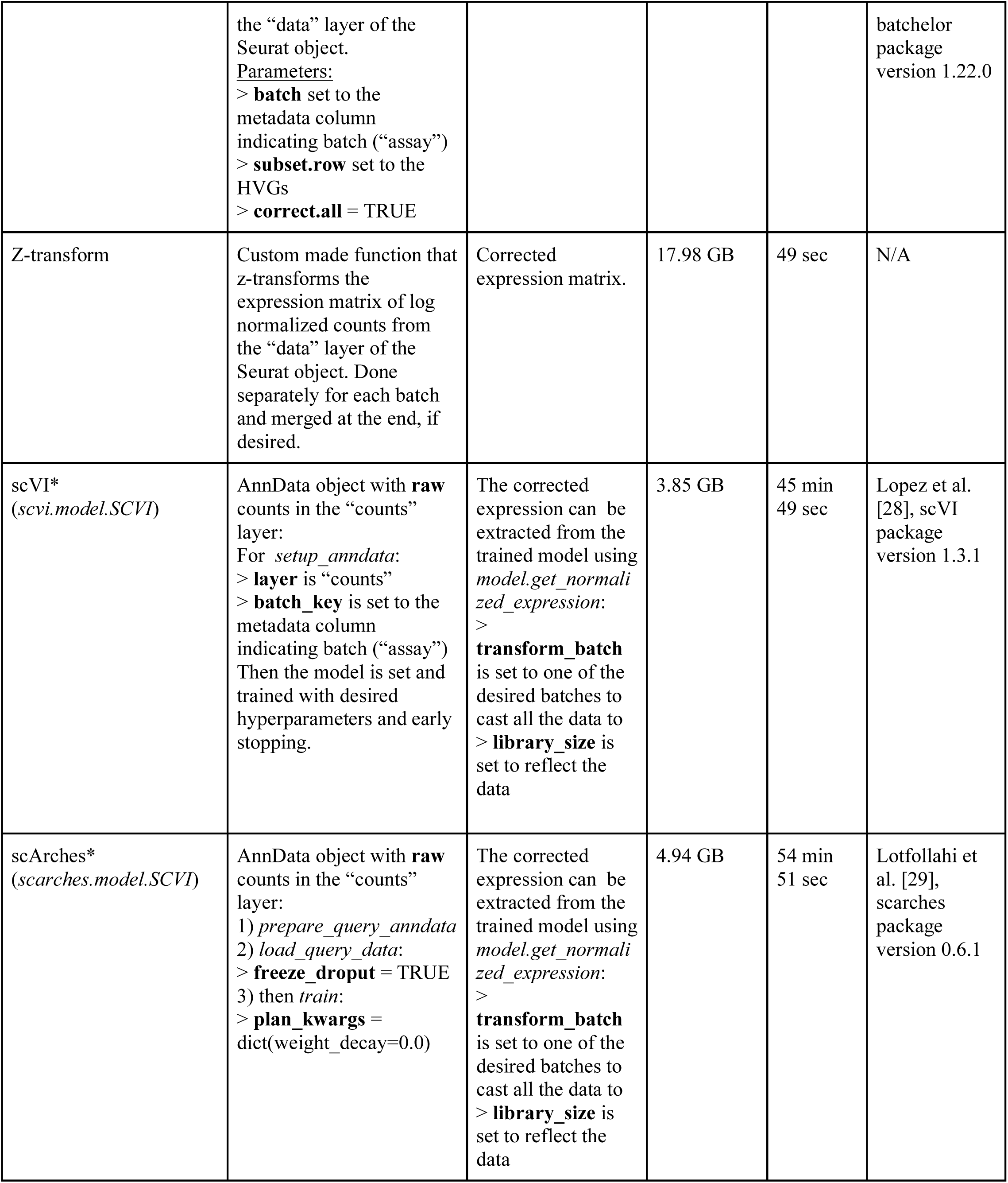
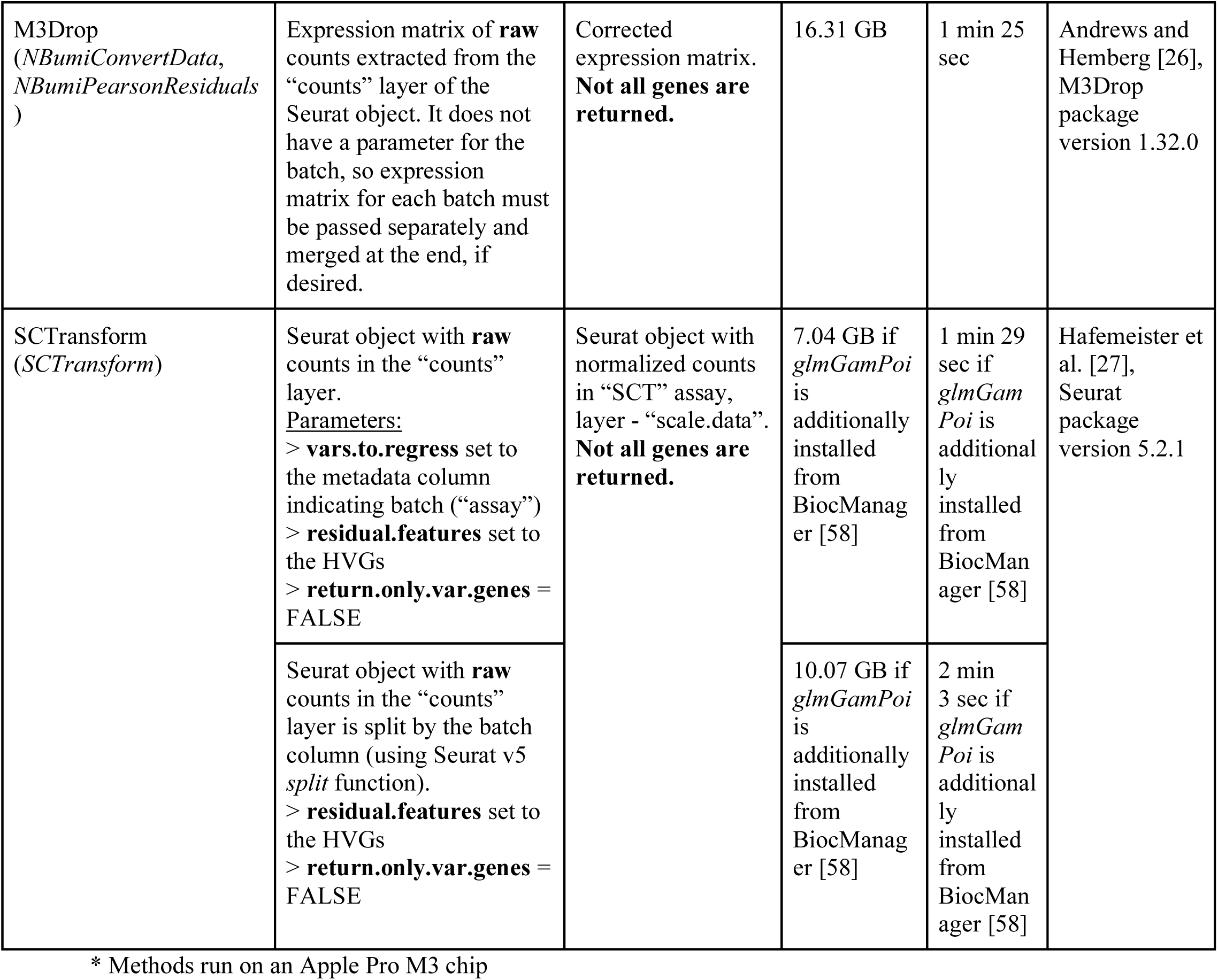
Description of batch correction methods used in this study.

#### SCTransform-regression

Normalization was performed using sctransform (v0.4.1) [27] implementation with default parameters on merged 3′ and 5′ data, with assay type (3′ vs. 5′) included as a covariate in the regression model.

#### SCTransform-single

sctransform (v0.4.1) [27] with default parameters was applied independently to 3′ and 5′ datasets, which were subsequently merged.

#### M3Drop

A depth-adjusted negative binomial (DANB) model was fit separately to 3′ and 5′ data using M3Drop (v1.32.0) [26]. Pearson residuals derived from fitted models were used as corrected expression values.

#### mnnCorrect

Log-normalized expression values from merged 3′ and 5′ datasets were corrected at the expression level using *mnnCorrect* from batchelor (v1.22.0) [25]. Due to computational constraints, this method was applied only to HVGs and consensus protocol-biased genes.

#### fastMNN

Normalization was performed as above using the *fastMNN* algorithm from batchelor (v1.22.0) [25].

#### ComBat

Log-normalized expression values from merged 3′ and 5′ datasets were corrected using the parametric implementation of *ComBat* in sva (v3.54.0) [23], which adjusts both the mean and scale of batch effects.

#### Limma

Log-normalized expression values from merged 3′ and 5′ datasets were corrected using the *removeBatchEffect* function from limma (v3.62.2) [24], which fits a linear model to each gene and removes systematic differences attributable to batch effects while preserving overall expression structure.

#### Z-transformation

3′ and 5′ datasets were independently normalized to the expression of the top 96 housekeeping (HK) genes as identified by Lin et al. [39]. For each cell, the average log-normalized expression of these genes was used to define a more stable “reference gene”. Expression values were then Z-transformed using the mean and standard deviation of this reference.

### scVI and scArches adaptation

In addition to the standard methods described above, we adapted scVI (v1.3.1) [28] for correcting 3′ vs 5′ data. Assay information was provided as a batch key during model training with no other covariates. Training was performed on the entire set of genes, and hyperparameters were set based on the developer’s recommendation. Normalized (decoded) gene expression for 3′ cells was then obtained using “get_normalized_expression” with *transform_batch* = 5′ assay, and with 10,000 as the library size.

Using scArches (v0.6.1) [29] we projected 3′ scRNA-seq data onto 5′ scRNA-seq space. We used the scVI autoencode model basis [28] and adapted the existing batch projection method to project 3′ into the latent space of 5′. Unlike scVI, we trained scArches using a leave-one-out cross-validation strategy where one entire dataset was excluded from training and used to validate model performance. Normalized (decoded) gene expression was then obtained using “get_normalized_expression” with *transform_batch* = 5′ assay, and with 10,000 as the library size. This reconstructed 5′ scRNA-seq data was compared to the matching real 5′ scRNA-seq data.

### Statistical performance measures

Since matched datasets were derived from the same tissue and subject but did not contain identical cells, we calculated the mean expression of every gene for each cell type within each dataset both before and after normalization. For each cell type, we computed Pearson correlation coefficient, Cosine similarity, Mean squared error (MSE), Euclidean distance, and Jensen-Shannon divergence (JSD) (philentropy v0.9.0) [40]. Performance was assessed using the percent change for Euclidean distance, Manhattan distance, and MSE. As for bounded metrics, such as Pearson correlation, Cosine similarity, and JSD, performance was assessed as percent change relative to the maximum change possible.

#### Controls

As M3Drop and SCTransform rescale each gene to a mean of zero and a variance of one, they were compared to the combined 3′ and 5′ data after log-normalization and scaling. All other normalization methods were compared to log-normalized expression.

### Comparing performance - percent and relative percent change

To compare the statistical performance of all correction methods, we calculated the percent change in each metric relative to the scores for the appropriate control (see Methods-Controls above). For bounded measures, we adapted the percent change calculated to account for the total possible improvement:

Bounded [-1,1] such as Correlation Coefficient and Cosine Similarity:

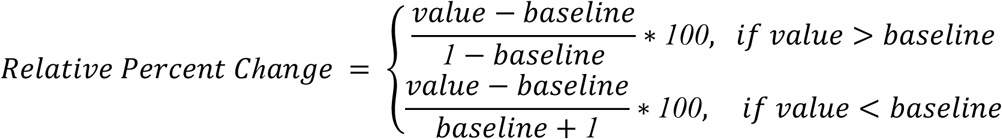

Bounded [0,1] such as JSD:

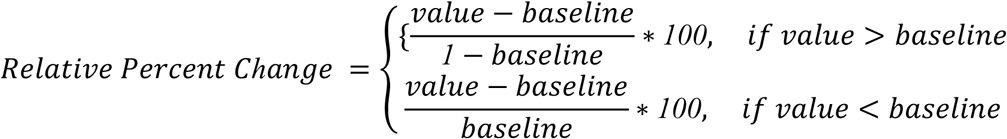

Unbounded such as Euclidean Distance, Manhattan Distance, and MSE:

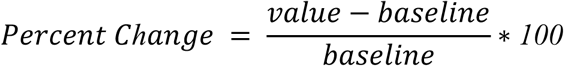

### UMAPs and mixing metrics

Corrected 3′ and 5′ expression matrices were merged and processed using the standard Seurat v5 workflow [37]. HVGs were obtained as described in “Data Preprocessing & Highly Variable Genes” section above. For methods other than M3Drop and SCTransform the data was scaled such that each gene had a mean of 0 and standard deviation of 1. Scaled-corrected data was subject to singular value decomposition for dimensionality reduction using Seurat (v5.1.0) to generate 40 principal components. Based on the proportion of variance explained by each component the top 20 principal components were selected for downstream analysis. UMAPs were generated using default parameters, and the ‘MixingMetric’ was calculated using Seurat. The neighbourhood size was set to 150 cells as a balance between looking at local and global scale.

### Marker-gene overlap

Cell-type-specific markers were identified within corrected 3′ and corrected 5′ scRNA-seq datasets independently using a Wilcoxon rank-sum test. Significance was determined as 5% FDR. Marker consistency was defined as the intersection of 3′ and 5′ marker genes divided by the minimum number of significant markers in 3′ or 5′. These Overlap Scores were averaged across subjects for each cell type. Uncorrected log-normalized counts were used as a reference.

### Biological use case construction

We selected three of the matching 3′ and 5′ datasets (Park thymus [31], Suo liver [33], Jardine bone marrow [32]) to generate synthetic use-case scenarios and evaluate top-performing correction methods, as these datasets contained similar yet distinguishable cell types that were also sufficiently abundant within each subject (Table 1). For each dataset, sparsity was introduced by subsetting 3′ and 5′ data such that a target abundant cell type was unique to each assay type (Table S5). DEGs were identified between the two target cell types after correction and compared with the ground truth DEGs (outlined below). Datasets with 100% overlap (original samples) were used as controls.

The ground truth DEGs were defined for each dataset by identifying consistent significant DEGs between the two chosen cell types in 3′ and 5′ samples and intersecting these results across all donors. Significance was determined as 5% FDR. To account for limited power, we defined true positives (TP), false positives (FP), true negatives (TN), and false negatives (FN) as follows (shown in Fig. 4B): TPs were significant in the ground truth and in the corrected comparison and were upregulated in the same cell type; FPs were any genes that were upregulated in the opposite cell types between the ground truth and the corrected values; FNs were genes upregulated in the same cell type and significant in the ground truth but not significant in the corrected values; TNs were genes upregulated in the same cell type but not significant in either the ground truth or the corrected values. These definitions were used to calculate Matthews correlation coefficient (MCC). We also calculated the total number of significantly up- and down-regulated genes identified by each method at 5% FDR.

### Computational demands

Run time and memory usage were evaluated using the Park dataset which comprised 9 subjects with an average of 4,415 cells per subject (total 39,709). Each correction method was run on a Digital Research Alliance of Canada (alliancecan.ca) high performance computing cluster node (Rorqual) with 2 x AMD EPYC 9654 (Zen 4) @ 2.40 GHz, 384MB cache L3. The exception was made for scVI and scArches which were run on a personal computer with an Apple M3 Pro chip.

## Results

### Exploration of protocol-biased genes between 10X Genomics 3′ and 5′ across 35 matched samples

We performed differential gene expression analysis on 35 donor samples with matched 10X 3′ and 5′ scRNA-seq data, drawn from six publicly available datasets spanning multiple tissues (Table 1) [31–36]. 3′ and 5′ data was compared for each subject separately to avoid any confounding by inter-individual differences. We identified between 609 and 2,598 genes consistently biased between platforms at different significance thresholds (1e^-5^, 1e^-50^, and 1e^-100^) (Fig. 1A).

**Fig. 1:**
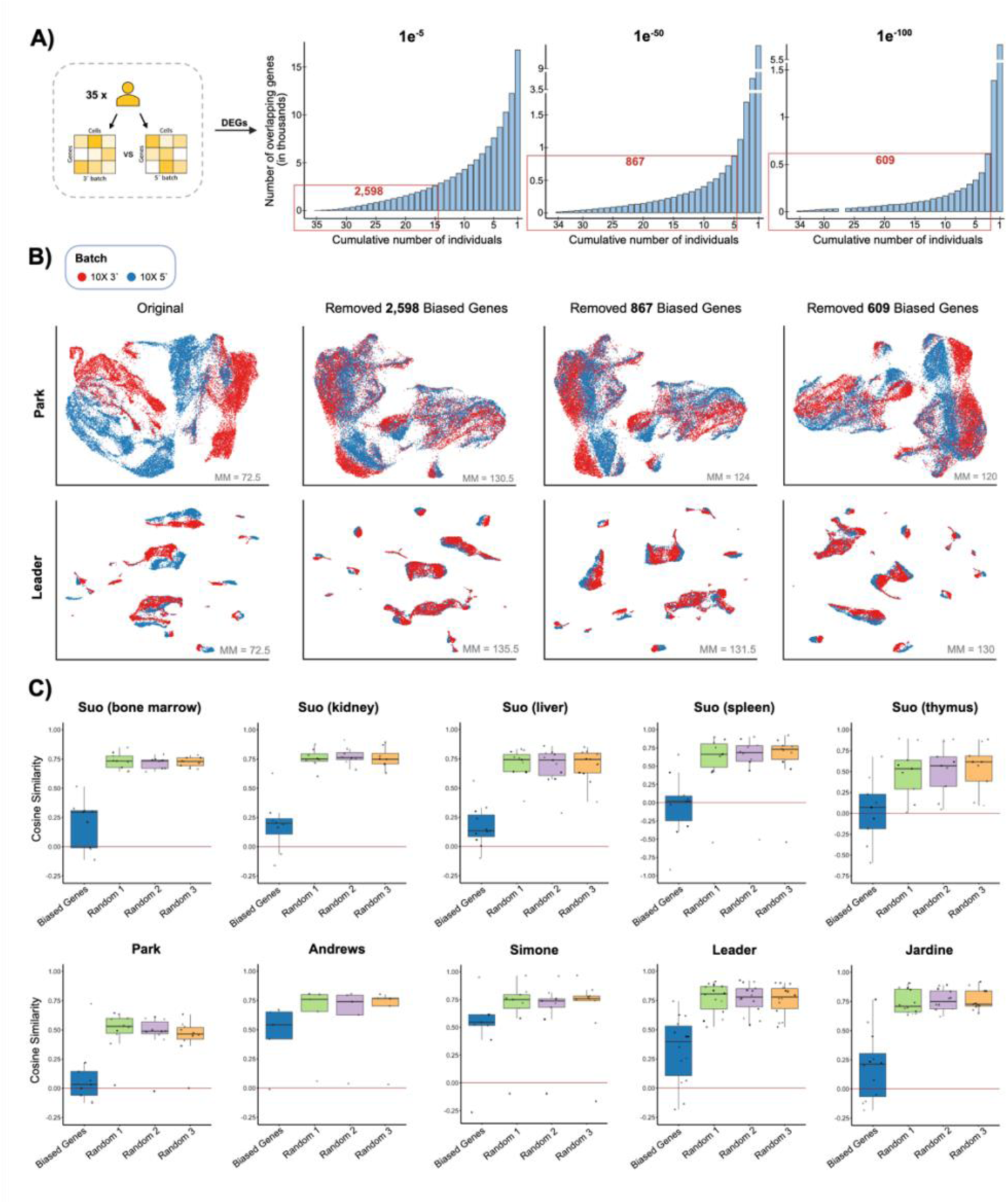
Exploration of differentially expressed genes (DEGs) between 10X 3′ and 10X 5′ protocols across 35 matched samples. 35 pre-processed and 3′/5′ matched scRNA-seq samples from 6 datasets of different tissues were obtained from the CellXGene platform and analyzed in R (v4.4.1) using Seurat (v5.1.0). (A) Cumulative distribution of overlapping DEGs across individuals at three significance thresholds (1e⁻⁵, 1e⁻⁵⁰, 1e⁻¹⁰⁰). Bars indicate the number of shared DEGs as more individuals are considered cumulatively; red labels show the total DEGs chosen at each threshold for further investigation. (B) UMAP visualizations from representative datasets (Park and Leader) showing cells from 10X 3′ (red) and 10X 5′ (blue) protocols. The first column shows the original data, while subsequent columns display cell distributions after removal of Biased Genes (DEGs corresponding to the chosen number in red from the Panel A). For each plot, the value of the Seurat Mixing Metrics is indicated in grey at the bottom right corner, with higher values representing a better batch mixing. (C) Boxplots of Cosine similarity between average gene expression profiles of 3′ and 5′ data, comparing scores for the 867-gene list (1e⁻⁵⁰ threshold) to three replicates of randomly selected gene sets of equal size across five representative datasets. Cell types are shown as different shapes for noticing trends.

By further removing the three sets of biased genes, we observed an improvement in the overlap between 3′ and 5′ data in the lower dimensional space (Fig. 1B). UMAPs showed clear batch separation in the original data that was largely eliminated when biased genes were excluded before preprocessing (Fig. 1B, Fig. S2). In addition, the ‘MixingMetric’ [38] showed a striking reduction upon removing as few as 609 genes, with a smaller decrease as additional genes were removed. We selected the 867-gene set, hereafter referred to as the *biased genes*, for downstream analysis as it provided a balanced compromise - smaller gene sets resulted in reduced alignment of 3′ and 5′ data in several datasets, while larger sets offered little additional improvement while removing substantially more genes (Fig. 1B, Fig. S2). For the biased genes, cosine similarity of average expression between 3′ and 5′ was below 0.25 in six of ten datasets and substantially lower than for other genes across all datasets, demonstrating a strong technical effect only on biased genes (Fig. 1C). In contrast, other genes typically had a cosine similarity of 0.5-0.75, reflecting high reproducibility across 3′ and 5′ technologies.

These results indicate that a relatively small subset of genes accounts for the majority of the expression differences between 10X 3′ and 5′ chemistries, while the remainder of the transcriptome appears largely consistent across protocols.

### Benchmarking of ten common scRNA-seq batch correction methods for correcting 10X 3′/5′ protocol bias

We compared the performance of 10 well-known normalization methods (Table 2) [23–29] in their ability to correct the gene-expression values of 3′ and 5′ scRNA-seq data while preserving the biological variation (Fig. 2A). To mitigate confounding effects from inter-individual genetic heterogeneity, all normalization methods were applied independently to the matching 3′ and 5′ samples for each of the 35 subjects. Samples with substantial differences in cell-type frequency between 3′ and 5′ data were excluded to avoid biases from incomplete dissociation. An exception was made for the Andrews and Suo (thymus) datasets to introduce a more challenging scenario while ensuring sufficient data was included.

**Fig. 2:**
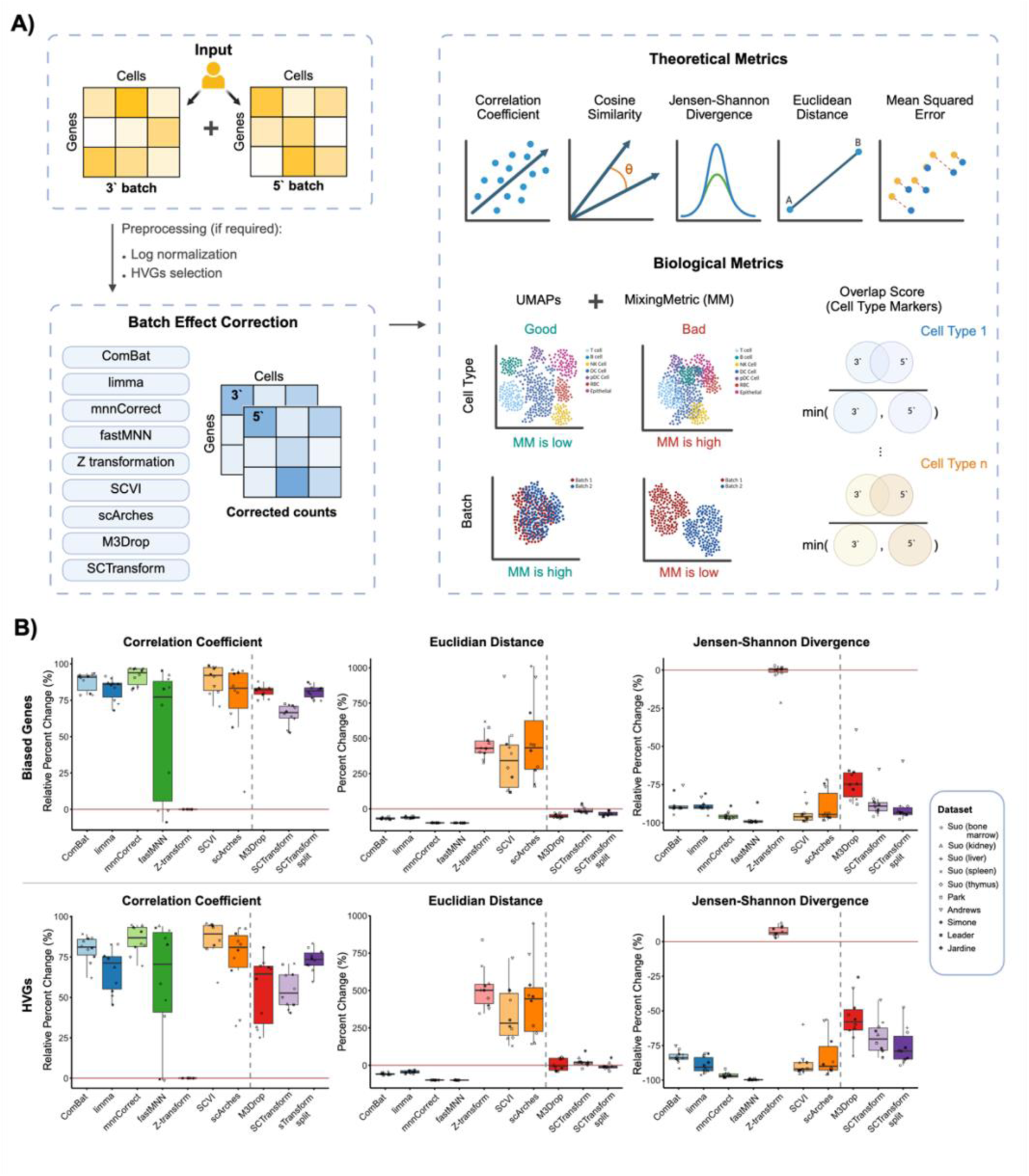
Benchmarking of ten common scRNA-seq batch correction methods for correcting 10X 3‘/5’ protocol bias. 35 matched samples from 6 datasets were independently integrated using each technique. (A) Benchmarking workflow. (B) Boxplot of three representative metrics comparing the ten correction techniques across all datasets (shown as different shapes), with zero (red line) indicating the respective baseline. The dashed gray line indicates that the baseline used for ComBat – scArches is different from that of M3Drop and scTransform (log-normalized vs long-normalized-scaled counts respectively). Higher correlation coefficients and lower values for other metrics denote better integration performance. Correlation coefficient and JSD results are shown as relative percent changes; Euclidean distance results are shown as percent changes. Top shows the metrics for “Bias Genes” from Figure 1, bottom shows the metrics for the 3,000 Highly Variable Genes. Created with BioRender.com.

### Quantitative measures of corrected 3′ vs 5′ similarity

Firstly, the performance was assessed using statistical metrics by comparing cell-type-specific expression of the previously identified biased genes and 3,000 highly variable genes (HVGs) between the corrected 3′ and 5′ scRNA-seq data relative to an appropriate reference (Fig. 2B, Fig. S3). For the biased genes, we found that ComBat [23] and mnnCorrect [25] consistently yielded the largest improvements over the uncorrected baseline, they increased the Pearson correlation between 3′ and 5′ expression, and reduced JSD and Euclidean distance. Whereas our housekeeping-gene-based Z-transformation performed the worst with no evidence of improving 3′ and 5′ concordance.

Other methods had differing results depending on the metric used. FastMNN [25] performed similarly using Euclidean distance and JSD but had variable performance using Pearson correlation. Limma [24] performed similarly to ComBat except for weaker performance on the correlation. The machine-learning approaches, scVI [28], and scArches [29], were comparable in performance to the others using the correlation and JSD but substantially increased the Euclidean distance between 3′ and 5′ vs baseline.

In contrast to the other methods, the Pearson-residual-based approaches, M3Drop [26] and SCTransform [27], showed substantially better performance on the biased gene set than on the HVGs, particularly when considering the Euclidean distance and, for SCTransform, MSE (Fig. S3A). Somewhat surprisingly, SCTransform run using its “split” option, which fits regression independently for each sample, substantially outperformed the regression-model approach used to correct for 3′ vs 5′ differences across all metrics and gene sets. Results using Cosine similarity and MSE were consistent with those for Pearson correlation and Euclidean distance respectively (Fig. S3).

### 3′ vs 5′ integration and cell-type marker conservation

In addition, we assessed the performance of each normalization method to align 3′ and 5′ datasets in lower dimensional space. Corrected expression data was merged, decomposed into 20 principal components, and visualized using UMAPs (Fig. 3A). The top performing methods were ComBat, fastMNN, and M3Drop which produced complete overlap of 5′ and 3′ scRNA-seq datasets without disrupting cell-type clustering. Whereas limma, Z-transformation, and SCTransform regression failed to integrate 3′ and 5′ datasets, having significantly lower ‘MixingMetric’ for 3′ vs 5′ labels (Fig. 3B). Similar to our observations above, the “split” approach to SCTransform consistently outperformed the regression model. In contrast, scArches effectively integrated 3′ and 5′, albeit at the cost of reduced cell-type distinctiveness. However, all methods, including scArches, preserved cell-type distinctions sufficiently to maintain a ‘MixingMetric’ of zeroes (Fig. S5).

**Fig. 3:**
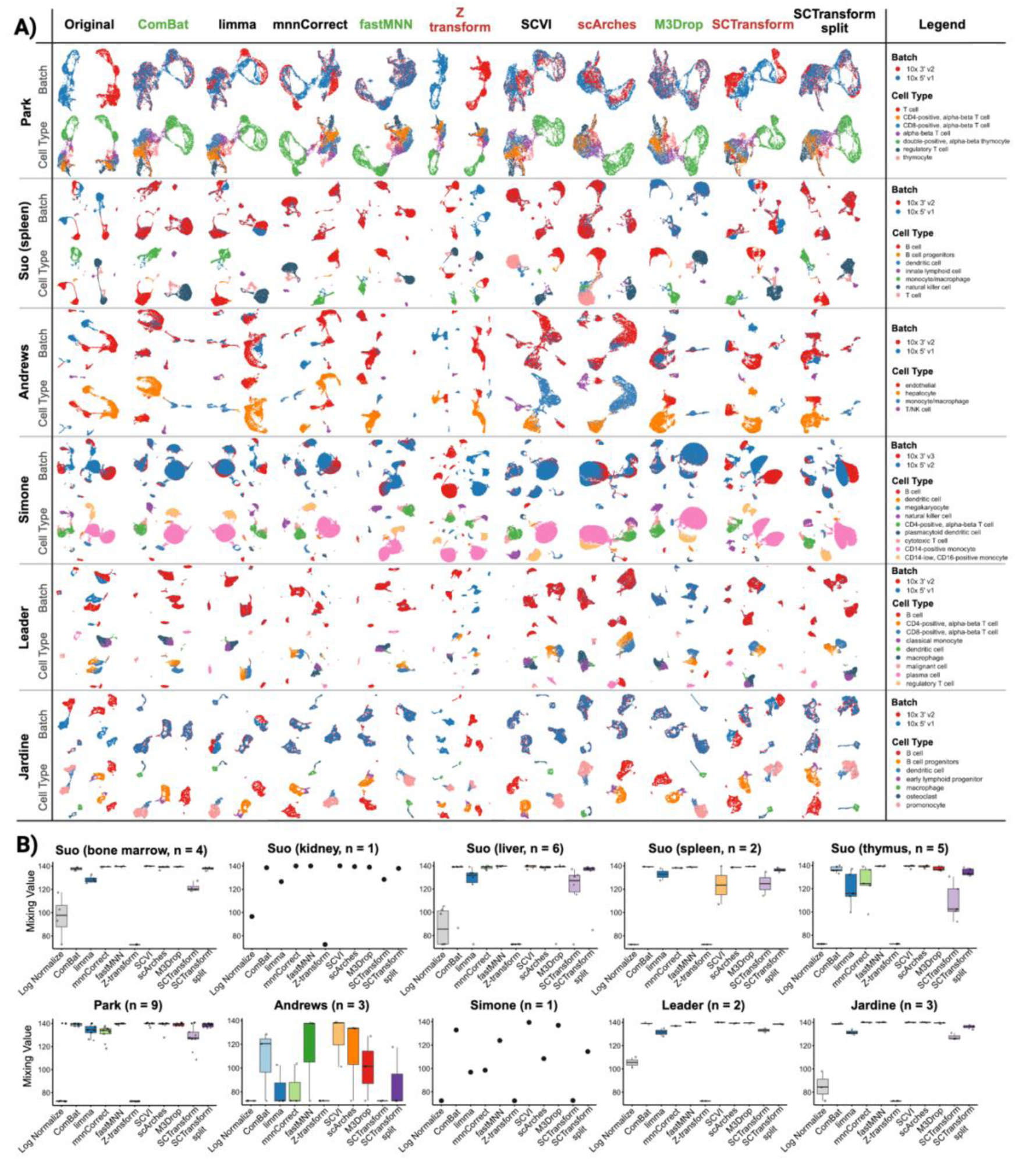
Evaluation of ten common scRNA-seq batch correction methods for preserving cell type structure after 10X 3′/5′ correction. 35 matched samples from 6 datasets were independently integrated using each technique based on the workflow shown in Figure 2A. (A) UMAP visualizations following different batch correction methods, using log-normalized counts as a reference. Cells are colored by sequencing protocol (top) and by cell type (bottom – as identified by the authors). Only representative plots of datasets and one individual per dataset are shown. The best techniques are indicated with green names while the worst - with red. (B) Boxplots showing batch Seurat Mixing Metrics results for all datasets. Shapes represent different individuals. Higher values indicate better mixing of cells across batches.

We also considered the consistency of cell-type-specific marker gene identification between 3′ and 5′ data (Fig. S4). While fastMNN, scVI and scArches achieved the highest marker overlap, this was likely a result of increasing the number of marker genes 6-10-fold to 7,000-30,000 for some cell types. Thus, it was unclear whether any methods truly improved the consistency of cell-type markers.

### Computational cost to correcting 3′ and 5′ gene expression

Most normalizations scaled well, requiring < 20 Gb of RAM, and completed in < 5 minutes of runtime on 2 x AMD EPYC 9654 (Zen 4) CPUs at 2.40 GHz, 384MB cache L3. The exception was mnnCorrect which required 30.55 Gb of RAM and 51 min of runtime for the Park dataset [31] comprising 9 subjects with an average of 4,415 cells per subject and ∼ 40,000 cells total (Table 2).

### Applying batch correction methods to integrate partially overlapping 3′ and 5′ datasets

Finally, we simulated a real-world application scenario - correcting datasets with incomplete cell-type overlap where one dataset was collected with 3′ and the other with 5′. We introduced artificial sparsity into three selected datasets of different tissues (thymus, liver, and bone marrow) such that only ∼25% of cell types were shared among 3′ and 5′ samples, with the remaining cell types being unique to each batch (Fig. 4A). We also considered a scenario with complete (100%) overlap, using the original samples as controls. We focused our evaluation on the well-performing normalization methods identified in earlier analyses, assessing their ability to recover correct cell-type markers between two selected populations. Limma was chosen in place of ComBat, as an algorithmically similar linear method, since ComBat does not handle large cell-type discrepancies between integrated datasets well [25,41].

**Fig. 4:**
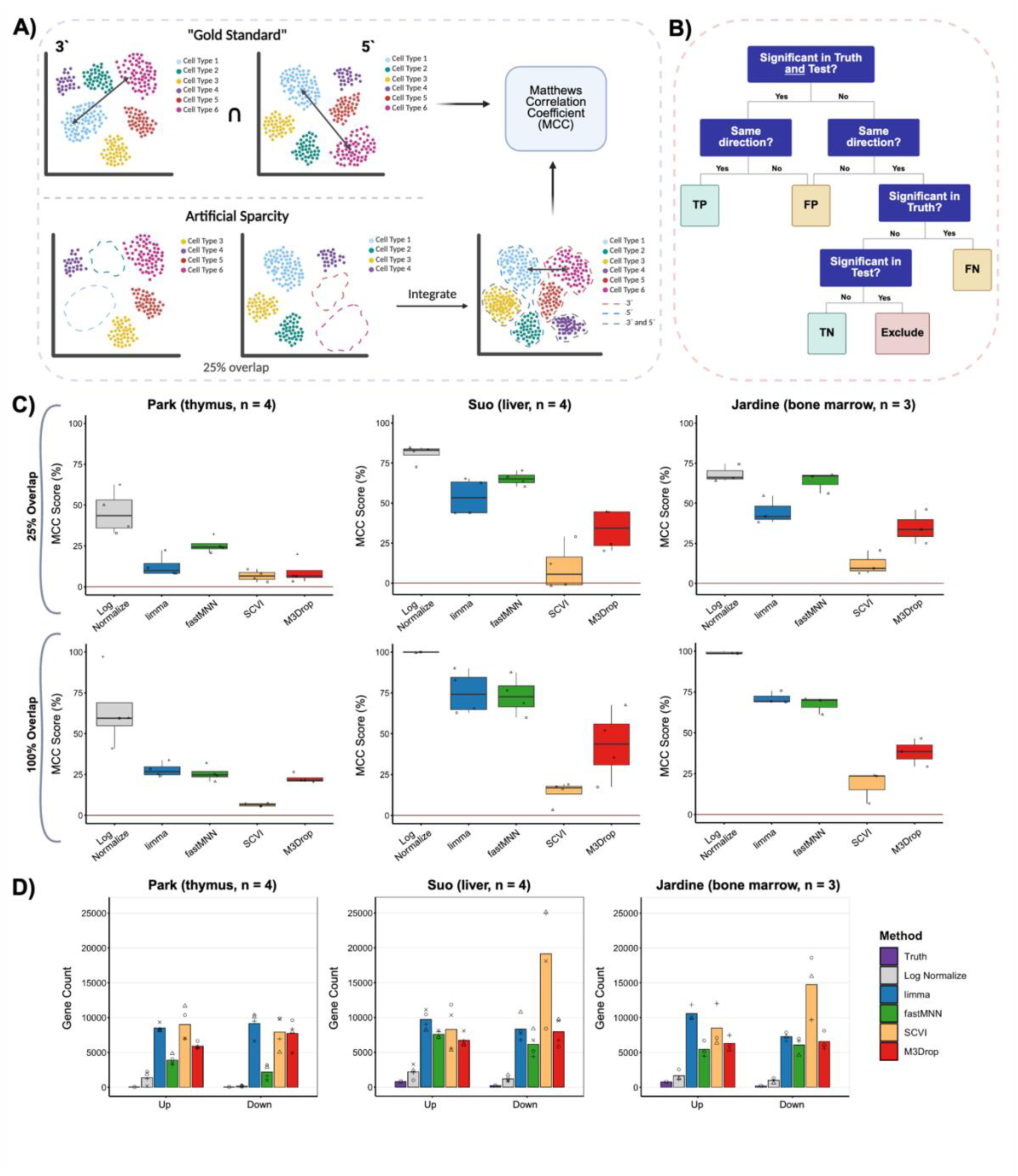
Evaluation of select batch correction methods in their ability to recover correct cell type markers. For each donor, selected batch correction approaches (limma, fastMNN, SCVI, and M3Drop) were evaluated for their ability to recover cell-type specific differential expression after 3′/5′ library integration in three selected datasets - Park (n = 4 donors), Suo liver (n = 4 donors), and Jardine (n = 3 donors). Log Normalization was used as an unintegrated reference. (A) Workflow diagram. The “Gold Standard” DEG set was defined as the intersection of genes differentially expressed between the two chosen cell types within 3′/5′ separately, followed by their intersection. Further, an intersection across individuals was performed to create one standard. Per individual integration performance on the artificially sparse dataset (in which 25% of cells overlapped between 3′ and 5′, but the compared cell types, along with some other cell types, were dropped out in opposite protocols) was quantified using Matthews correlation coefficient (MCC). (B) MCC was calculated based on custom TP/FP/FN/TN definitions. (C) Boxplots show the distribution of MCC scores across individuals (shown as different shapes) for chosen batch correction methods and datasets; higher MCC indicates greater agreement with the gold standard. Results from the artificially sparsed datasets are shown in the top panels, and results from the full (unsparced) datasets are shown in the bottom panels. (D) Bar plots show the number of significantly up- and down- regulated identified genes based on a set threshold after correction (limma, fastMNN, SCVI, M3Drop) using log normalization and “Gold Standard” Truth as reference. Created with BioRender.com.

To evaluate performance in these scenarios we identified differentially expressed (DE) genes between two cell types with high frequency in all subjects and moderate biological distinctiveness. Ground truth was generated for the selected cell-type pairs in each dataset by comparing these cell types within the 3′ and 5′ data and selecting those genes that were consistently DE in both assays across all subjects (Fig. 4A). Accuracy of each method was quantified using Matthews correlation coefficient (MCC), comparing the markers recovered after the correction to the ground truth DE genes using a power-agnostic classification (Fig. 4B). Surprisingly, uncorrected log-normalized expression yielded the highest MCC scores in both the sparse (25% overlap) and full-overlap settings (Fig. 4C), suggesting that in this context, batch correction may not enhance, and can even impair, marker gene detection accuracy. Among corrected methods, fastMNN stood out for its consistent performance even in the 25% overlap scenario, achieving MCC scores close to the reference. Moreover, compared to other correction techniques, fastMNN produced a DEG list with minimal inflation of gene counts, indicating a better balance between batch effect removal and specificity of differential signal recovery (Fig. 4D).

This poor performance was not attributable to the selected ground-truth marker genes lacking differences between 3′ and 5′ assays, as in all three cases at least 10% of the ground truth markers overlapped with our biased genes set, that intrinsically has large differences between 3′ and 5′ assays (Fig. 5A). Indeed, inspection of some classic cell-type markers for our chosen cell types, which were found among the ground truth genes, showed clear differences between 3′ and 5′ (Fig. 5B).

**Fig. 5:**
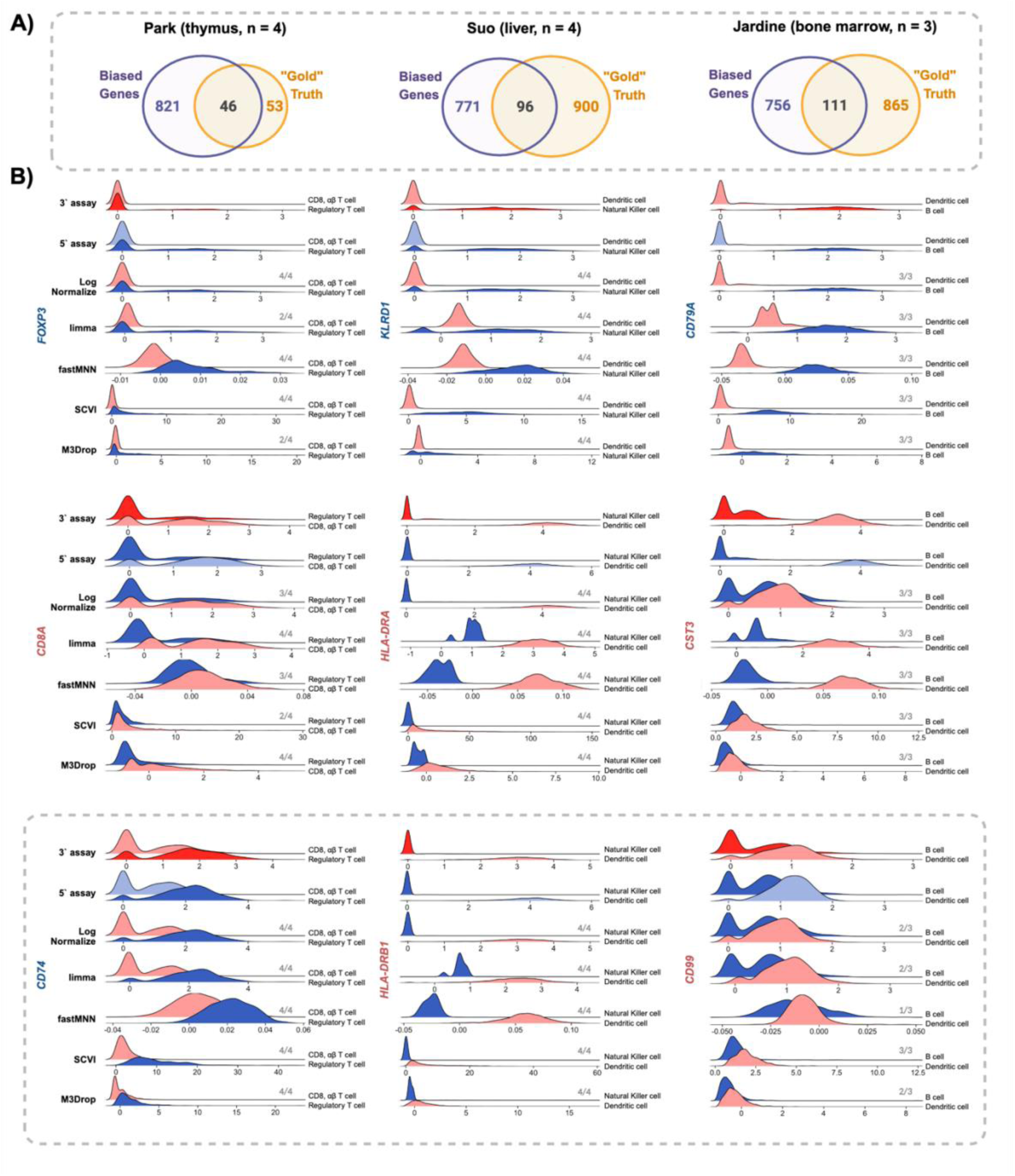
Evaluation of cell marker gene expression for select integration methods. Park (n = 4 donors), Suo liver (n = 4 donors), and Jardine (n = 3 donors) datasets were preprocessed as described in Figure 4 (Panel A). (A) Overlap between Biased Genes (genes consistently differentially expressed between 3′ and 5′ assays within each dataset, refer to Figure 1) and the “Gold” Truth marker gene set (genes consistently differentially expressed between the two target cell types across individuals, refer to Figure 4) for the selected datasets. Numbers within Venn diagrams indicate the intersection size between protocol-biased genes and gold-standard markers, with outer counts showing the total number of biased genes (purple) and gold-standard genes (orange) identified in each dataset. (B) Ridge plots illustrating the gene expression distributions for representative marker and protocol-biased genes after variable normalization techniques. For each dataset, the first two gene groups show true marker genes—one for each of the two cell types being compared. Within each gene group, the first two rows show expression from the unsparced 3′ (red) and 5′ (blue) assays, using log-normalized counts. Subsequent rows show the same genes after artificial sparsification (as described in Figure 4) and application of downstream normalization methods (log-normalization, limma, fastMNN, SCVI, and M3Drop), again comparing the two cell types (red = cells retained in 3′; blue = cells retained in 5′). The final gene group (outlined with a grey dashed box) displays an example overlap gene, i.e., a gene present in both the protocol-biased list and the gold-truth marker set, demonstrating how such genes can retain protocol-driven expression differences even after correction. The gene name colour represents which cell type the gene is the canonical marker for (red – cell type retained in 3′; blue – in 5′). The grey numbers for each Ridge Plot show how many individuals had this gene in their marker gene list for the respective cell type after correction.

While correction methods did significantly alter the expression of these genes, they failed to accurately remove the specific bias introduced by 3′ vs 5′ technologies. Notably, fastMNN and scVI generally smoothed and imputed expression values in addition to correcting them, which successfully preserved DE patterns for genes with clear differences, but which sometimes led to over-smoothing in genes with more overlapping distributions such as CD99. fastMNN also uniquely compressed the range of gene-expression values, with corrected values typically falling within [0, 0.1] range. Meanwhile, limma and M3Drop tended to alter zero values in a manner that contradicted the correct DE pattern, leading to a loss of significance in genes such as FOXP3 and KLRD1. Notably, only limma consistently preserved bimodality of expression when present, as observed for CTS3 and CD99 genes.

These findings highlight the challenge of correcting protocol-specific effects without compromising biological variation. Although 10X 3′ and 5′ protocols introduce detectable expression biases, these differences are largely confined to a small subset of genes. Across matched samples from diverse tissues, most batch correction methods performed comparably on statistical benchmarks, with fastMNN, mnnCorrect, and SCTransform (split) showing consistently strong alignment of expression profiles. However, when evaluated in a biologically relevant context, the performance shifted. Methods like fastMNN maintained robust marker recovery, while others that performed relatively well in statistical metrics, such as scVI or M3Drop, showed inflated gene detection or distorted expression dynamics. Importantly, we found that simply excluding the small set of 867 protocol-biased genes prior to analysis substantially improved downstream alignment and may provide a more reliable alternative to complex correction workflows in certain scenarios.

## Discussion

Both 3′ and 5′ scRNA-seq chemistries are widely used, with the 5′ protocol commonly employed for immune repertoire analysis due to its ability to capture recombined TCR and BCR sequences [11,42], while 3′ protocol remains prevalent in large single-cell atlases due to its cost efficiency and historical use [1,6,43,44]. As a result, publicly available scRNA-seq datasets increasingly comprise a mixture of 3′ and 5′ chemistries, creating both opportunities and challenges for cross-protocol comparisons and meta-analyses. Most existing batch correction methods only align datasets in a reduced-dimensional space, rather than correcting gene-level expression values directly [15,17]. Clear guidance on whether - and how - gene expression values from 3′ and 5′ protocols can be meaningfully compared is lacking. Our study addresses this gap by characterizing protocol-specific expression differences and evaluating commonly used normalization strategies in biologically realistic settings.

Interestingly, the majority of protocol-induced variation was driven by a relatively small subset of genes (Fig. 1). This observation is consistent with the underlying chemistry of 3′ and 5′ scRNA-seq, where protocol-specific effects arise primarily from variation in capture and amplification efficiency across each transcript [45,46]. Unlike comparisons between scRNA-seq and snRNA-seq - where biological compartmentalization, nuclear retention, and cytoplasmic regulation introduce broad and confounded effects [19] - the 3′/5′ differences are isolated to a narrowly defined technical perturbation.

### Performance of gene-level batch correction methods across statistical benchmarks

Across multiple statistical metrics and datasets, fastMNN, mnnCorrect, and ComBat consistently improved agreement between matched 3′ and 5′ expression profiles as measured by statistical similarity (Fig. 2). These findings align with previous benchmarking studies showing that mutual nearest neighbor-based approaches and linear batch correction methods perform robustly in settings with moderate batch effects and shared cell populations [15,25,47]. While mnnCorrect often achieved slightly stronger alignment than fastMNN, it did so at a substantially higher computational cost, both in memory usage and runtime, making it prohibitive for the use in large-scale datasets or atlasing efforts. ComBat also performed well on statistical metrics, but its reliance on global linear adjustments limits its applicability in scenarios with strong cell-type imbalance or incomplete overlap between batches [15,23]. Notably, the Andrews dataset [34], which had more divergent cell-type frequencies between 3′ and 5′ data, was the most difficult for the methods to correct, and only fastMNN and scVI were able to fully mix the two batches for this dataset (Fig. 3).

Interestingly, we observed a consistent difference in the behaviour of SCTransform variants (Figs. 2 and 3). Removing 3′ vs 5′ differences using a negative binomial regression model consistently underperformed compared to applying SCTransform separately to the 3′ and 5′ data, without any explicit attempt to remove the differences between the assays. A likely explanation for this observation is that SCTransform fits a regression model to a subset of genes and then interpolates regression coefficients across genes with similar baseline expression levels. We showed that 3′ vs 5′ differences are gene-specific and not consistent across genes of the same expression level, which would prevent SCTransform from accurately interpolating parameters of an appropriate correction. In contrast, when SCTransform is applied separately to each protocol, the difference in baseline expression levels of the protocol-biased genes is regressed to zero in each case, effectively eliminating the protocol bias.

### Statistical alignment does not necessarily translate to biological accuracy

When we further evaluated the normalization methods in a biologically relevant setting, involving comparisons between cell types present only in 5′ datasets with those present exclusively in 3′ datasets, a different pattern emerged (Figs. 4 and 5). In this scenario, uncorrected log-normalized expression consistently achieved the highest agreement in terms of differential expression between such cell types. Among correction methods, fastMNN remained comparatively robust but still underperformed relative to the uncorrected baseline.

This result highlights an important distinction between improving global similarity metrics and preserving biologically meaningful differences. Methods that performed well on distance- or divergence-based measures often identified substantially larger DEG sets, inflating both true positives and false positives (Fig. 4C). The machine learning methods, scVI and scArches, were particularly prone to inflating the number of DEGs, as one of their explicit goals is to denoise the data - a process that has previously been shown to inflate false-positive DEGs [48,49] (Fig. 4D). Our findings are consistent with prior observations that aggressive batch correction can obscure or distort differential expression signals, particularly when biological differences are subtle or partially confounded with batch structure [15,41].

### Implications: batch correction may be unnecessary for many 3′/5′ comparisons

Taken together, our results suggest that gene-level batch correction between 3′ and 5′ scRNA-seq data may be unnecessary - and potentially detrimental - for many common analytical tasks. Given that protocol-related differences are contained within a small gene set, a simpler strategy of excluding these biased genes may be sufficient to enable meaningful comparisons across protocols.

However, the sets of protocol biased genes we provide in Tables S6-S8 may not be exhaustive. Our definition is based on repeated Wilcoxon tests across donors using adjusted p value thresholds rather than effect size; some genes may be flagged due to small but consistent shifts that are statistically significant in large samples yet modest in magnitude, and conversely, genes with larger but donor-specific effects may be omitted. In addition, the bulk of our datasets were composed of immune-related cells, thus genes with low expression in these cell types may not have been detected as protocol-biased in our study. Expanding this work to include additional tissues would enable a more comprehensive identification of protocol-biased genes.

In addition, due to the design of this study we were able to control for many other sources of variation such as sequencing depth, tissue dissociation, and individual genetic difference, which would also confound attempts to compare expression between different published datasets. Similarly, we limit our analyses to intra-individual comparisons to isolate the effect of protocol bias rather than considering multi-individual between group comparisons, which are increasingly the focus of scRNA-seq studies [50–53]. Thus, it remains unclear whether statistically rigorous cross-dataset and/or multi-dataset comparisons are possible for scRNA-seq which may limit the potential of ongoing atlasing efforts [54].

Although our analysis focuses on differences between 10X Genomics 3′ and 5′ chemistries, our observations may extend to other closely related scRNA-seq technologies that differ only in enzyme formulations or RNA capture strategies, such as 10X v2 and v3. However, more substantially different technologies such as those using probe-based transcript quantification or different reaction volumes may have more pervasive and non-linear biases [55–57]. Prior work has shown that such technology-driven effects are often gene-specific and do not manifest as uniform global shifts across the transcriptome [18]. Thus, aggressive batch correction may distort quantitative gene expression and should be applied cautiously when batch effects are modest and biology is shared [15,47]. Since protocol-specific differences are technical in origin, it is at least theoretically possible to train a generalizable model to remove these differences. Our results suggest this remains an opportunity for the development of novel correction methods suited for such a task.

## AUTHOR CONTRIBUTIONS

VS: data curation, formal analysis, investigation, methodology, software, validation, visualization, conceptualization, writing - original draft, writing - revision and editing. TA: conceptualization, methodology, validation, supervision, writing - original draft, writing - revision and editing, funding acquisition. SMMH: conceptualization, funding acquisition, supervision, writing - revision and editing.

## CODE AVAILABILITY

The original code used for this analysis as well as the implementation of the existing R packages could be accessed through the GitHub (will be available upon publication).

## DATA AVAILABILITY

All datasets used during this study are publicly available on CellXGene. Individual datasets are listed in Table 1, along with CellXGene accession links.

## ACKNOWLEDGEMENTS

This research was enabled in part by support provided by Calcul Québec (calculquebec.ca), Compute Ontario (computeontario.ca), and the Digital Research Alliance of Canada (alliancecan.ca).

## AI USE STATEMENT

GenAI was not used for the generation of any information that has been presented in this study.

