## SupplementaryFigures for "Characterizing and Mitigating Protocol-Dependent Gene Expression Bias in 3′ and 5′ Single-Cell RNA Sequencing"

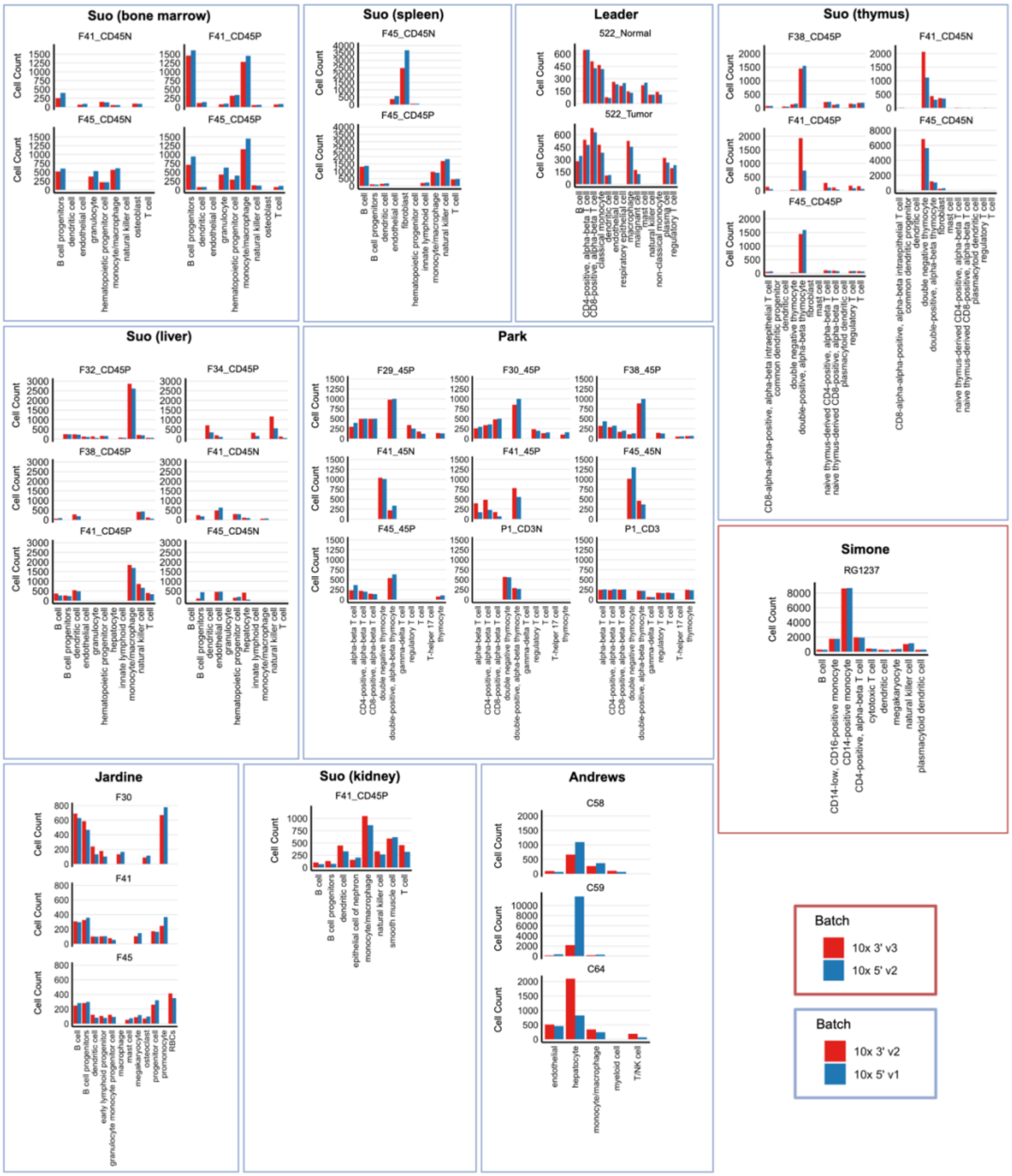

**Fig. S1: Quality control and assay composition of the analyzed datasets.** Bar plots show the number of cells per individual, stratified by sequencing assay/batch, across the selected datasets: Suo (bone marrow, spleen, liver, kidney, and thymus), Park, Andrews, Leader, Simone, and Jardine. Within each dataset, panels correspond to individual donors, and bars are colored by assay (red = 3′; blue = 5′). For five of the six datasets, the compared assays were 10x Genomics 3′ v2 and 10x Genomics 5′ v1. The Simone dataset instead compared 10x Genomics 3′ v3 and 10x Genomics 5′ v2. This distinction is indicated by the color of the bounding box surrounding each dataset panel and the accompanying legend (blue outline: 3′ v2 vs 5′ v1; red outline: 3′ v3 vs 5′ v2).

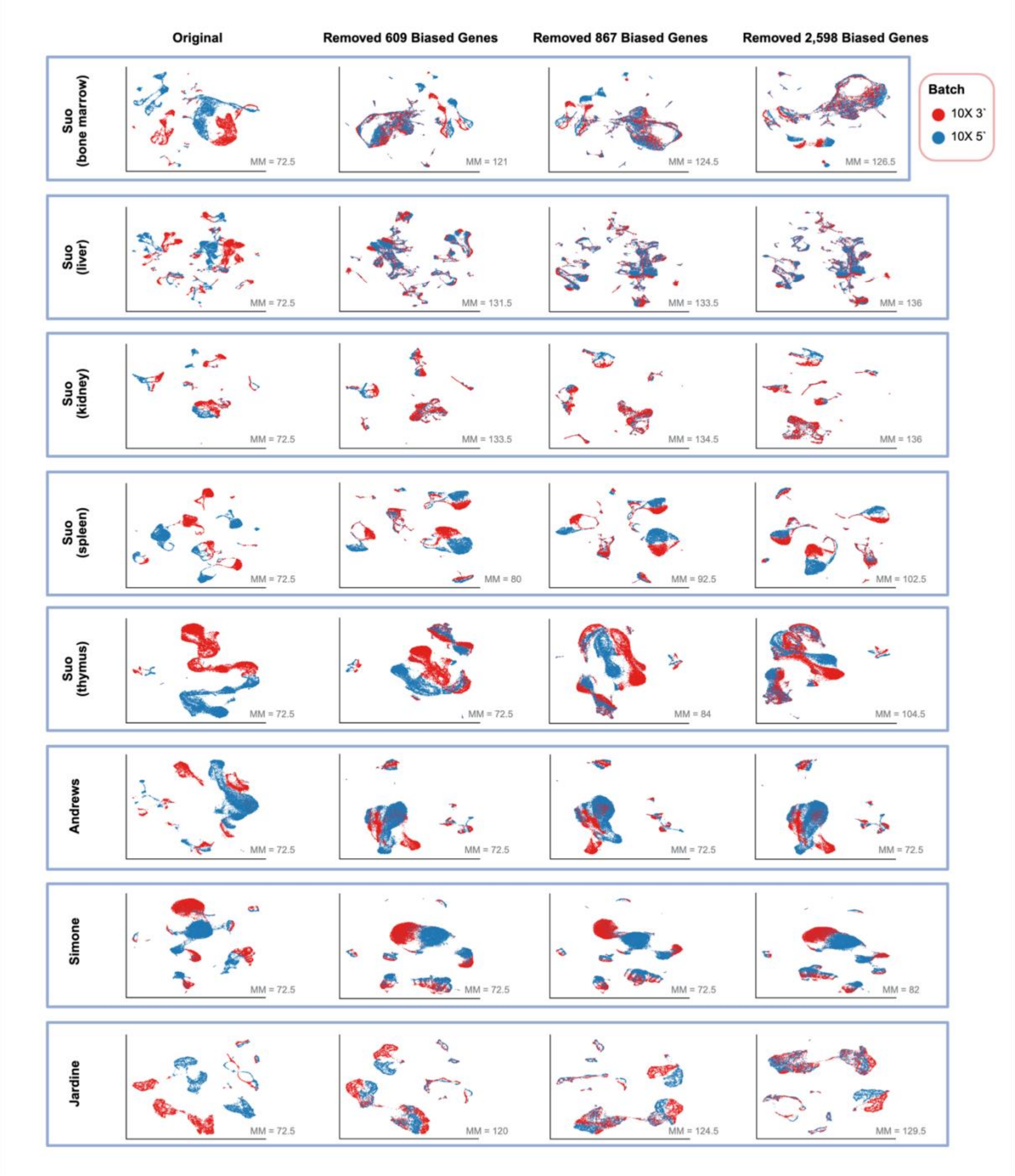


**Fig. S2: Removal of the protocol-biased gene sets across datasets.** UMAP visualizations for the remaining datasets—Suo (bone marrow, spleen, liver, kidney, and thymus), Andrews, Simone, and Jardine—showing cells generated using the 10X 3′ (red) and 10X 5′ (blue) protocols. For each dataset, four UMAP plots displays the original data as well as the same cells after removal of the 609, 867, and 2,598 gene sets (identified in Figure 1). For each plot, the value of the Seurat Mixing Metrics is indicated in grey at the bottom right corner, with higher values representing a better batch mixing.

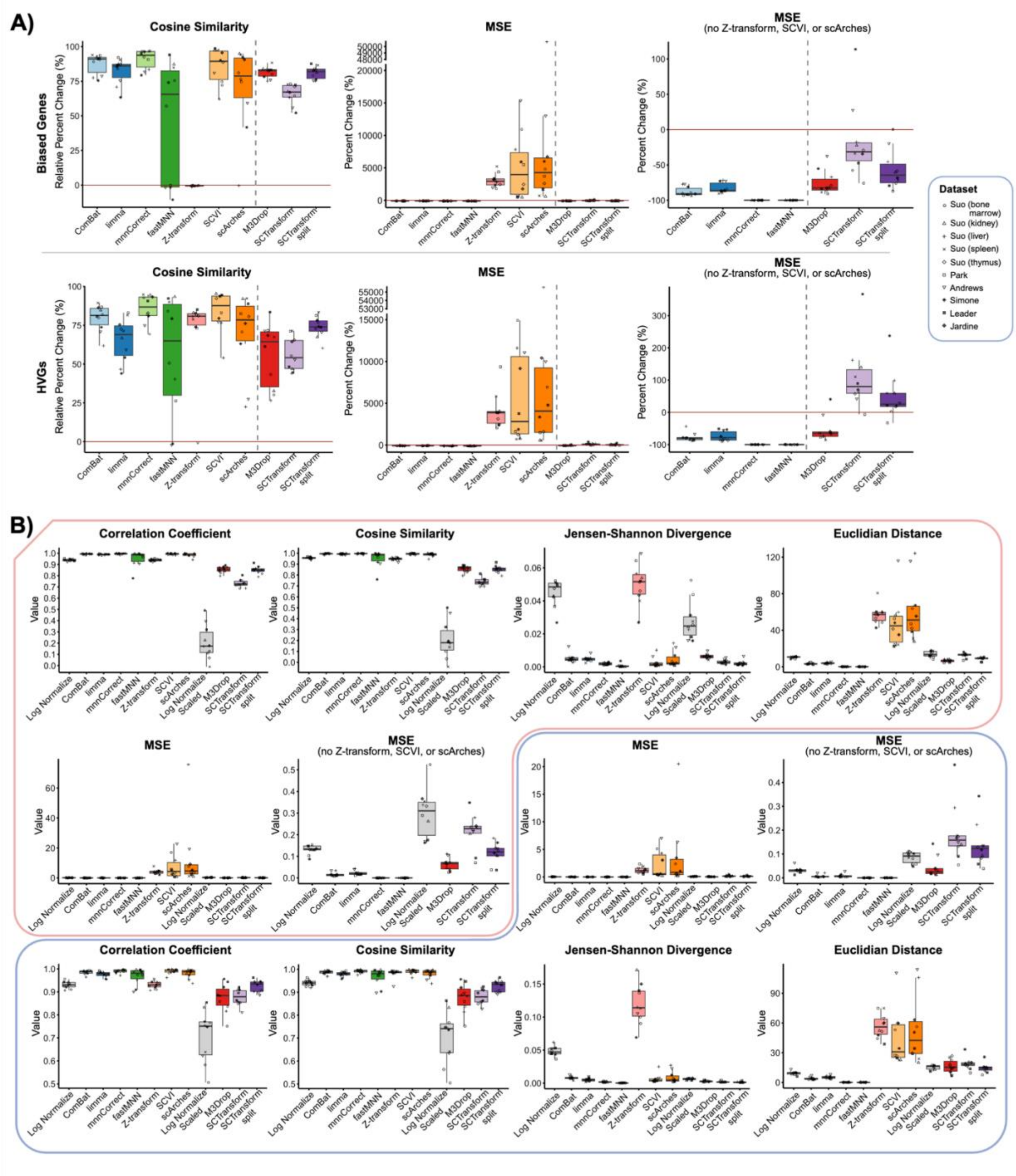


**Fig. S3: Extended results of benchmarking of ten common scRNA-seq batch correction methods for correcting 10X 3`/5` protocol bias.** 35 matched samples from 6 datasets were independently corrected using each technique. **(A)** Boxplot of Cosine Similarity, MSE, and MSE without some techniques (Z-transform, SCVI, and scArches) metrics comparing the integration techniques across all datasets (shown as different shapes), with zero (red line) indicating the respective baseline. The dashed gray line indicates that the baseline used for ComBat – scArches is different from that of M3Drop and scTransform (log-normalized vs long-normalized-scaled counts respectively). Higher correlation coefficients and lower values for other metrics denote better integration performance. Correlation coefficient and JSD results are shown as relative percent changes; Euclidean distance results are shown as percent changes. Top shows the metrics for “Bias Genes” from Figure 1, bottom shows the metrics for the 3,000 Highly Variable Genes. **(B)** Boxplots showing the raw metric values underlying the relative comparisons in (A), including Correlation Coefficient, Cosine Similarity, Jensen–Shannon Divergence, Euclidean Distance, MSE, and MSE excluding selected methods (Z-transform, scVI, and scArches). For each method, values cab be compared to their corresponding reference (Log Normalize or Log Normalize Scaled). The red outline highlights results computed using 867 “Biased Genes”, whereas the blue outline highlights results computed using the 3,000 HVGs.

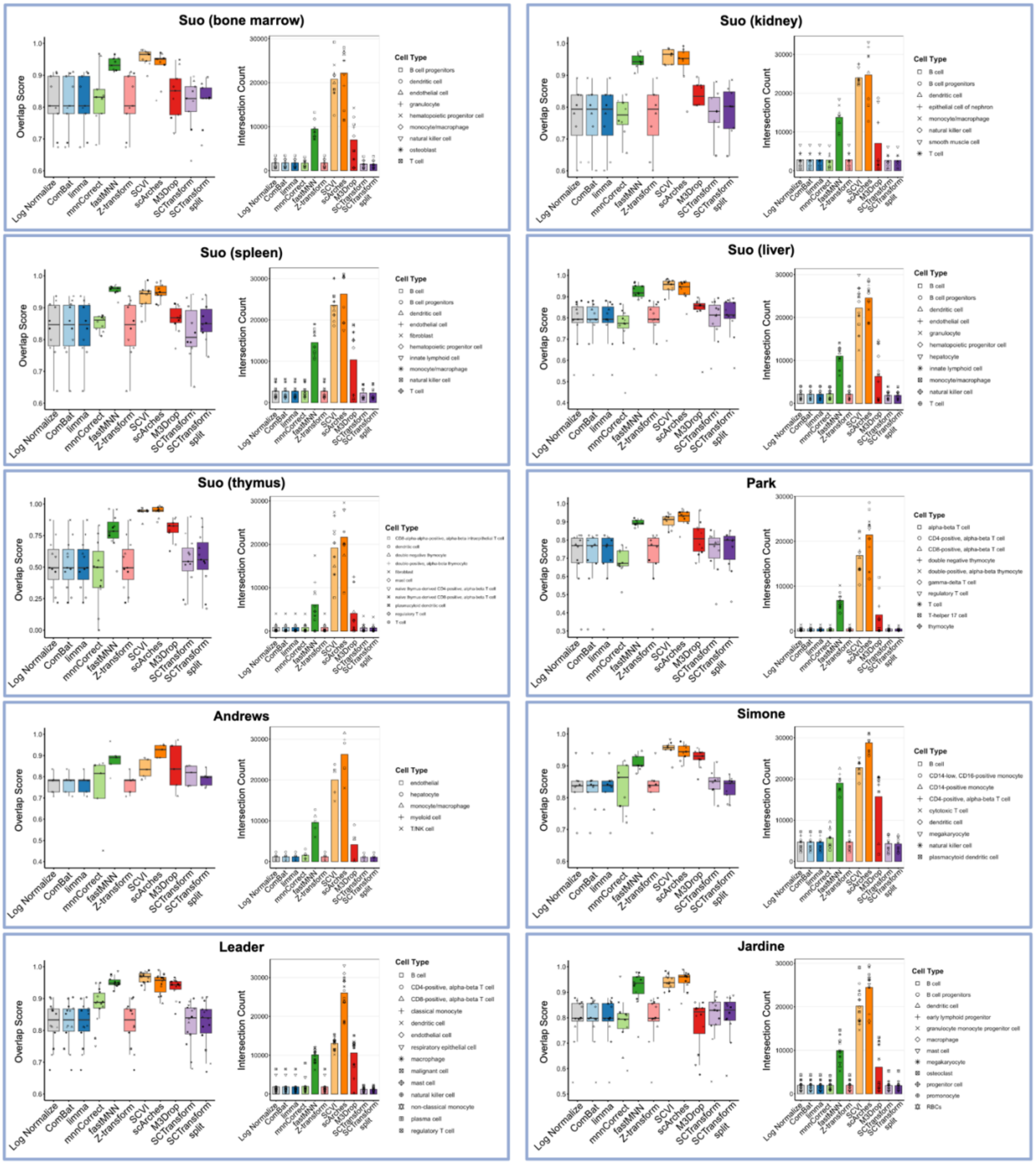


**Fig. S4: Evaluation of ten common scRNA-seq batch correction methods for preserving cell type markers.** Performance of ten commonly used scRNA-seq batch correction methods was evaluated across multiple datasets for their ability to preserve cell type–specific gene expression signatures after integration of 10x Genomics 3′ and 5′ protocols. Each dataset is enclosed by a blue outline and displayed as a paired set of panels. For each dataset, the **left panel** shows boxplots of the Overlap Score across different cell types (indicated by distinct point shapes). The Overlap Score quantifies the consistency of cell type marker identification between 10x 3′ and 10x 5′ data before and after batch correction, with higher values indicating greater agreement in marker recovery across protocols, as shown in Figure 2A. The **right panel** shows bar plots representing the number of overlapping cell type marker genes identified between 10x 3′ and 10x 5′ following correction. Log-normalized data serve as the reference baseline for comparison. Note that scTransform and MDrop automatically filter out some of the genes due to the QC reasons.

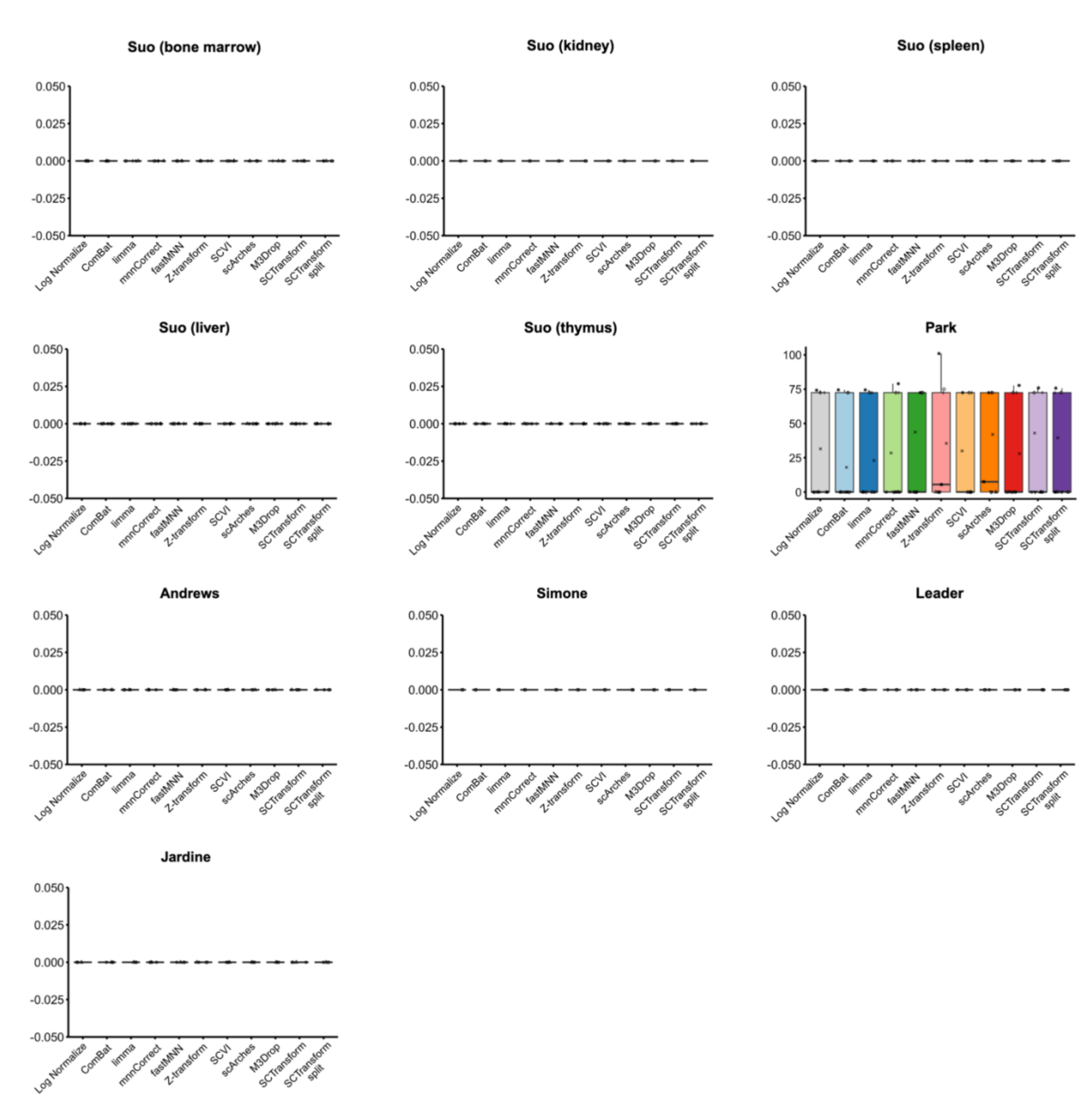


**Fig. S5: Evaluation of ten common scRNA-seq batch correction methods for preserving cell type structure after 10X 3′/5′ correction.** 35 matched samples from 6 datasets were independently integrated using each technique based on the workflow shown in Figure 2A. Boxplots show cell type Seurat Mixing Metrics results for all datasets. Shapes represent different individuals. Higher values indicate better mixing of cells across cell types, with lower values representing cell type integrity preservation.

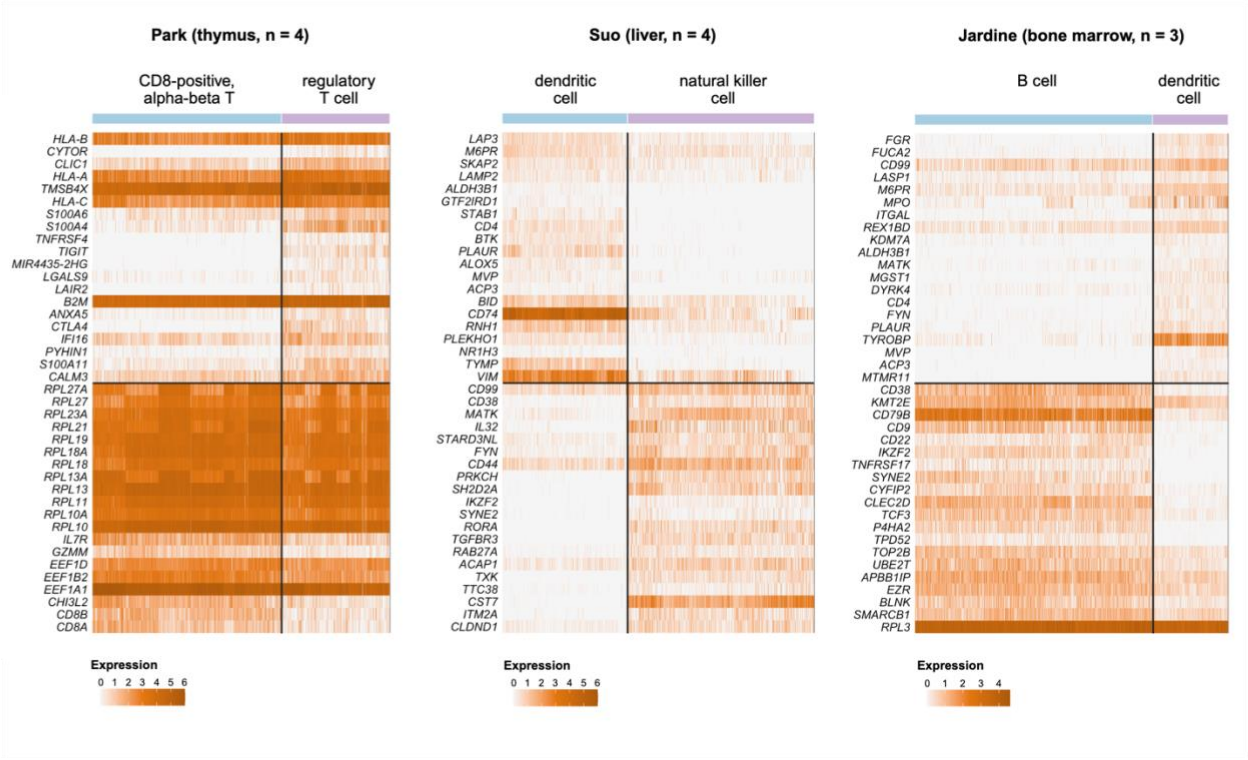


**Fig. S6: Validation of "Gold Standard" DEGs across three Use Case datasets and cell type comparisons.** Heatmaps display the expression patterns of "Gold Standard" DEGs identified between the two compared cell types in each dataset, created following the workflow described in Figure 4A. Results are shown for Park (thymus, n = 4 donors), Suo (liver, n = 4 donors), and Jardine (bone marrow, n = 3 donors). For each dataset, rows correspond to the top 20 up- and down- regulated genes included in the "Gold Standard" DEG set, and columns represent individual cells, grouped by cell type. Gene expression values are log-normalized within each dataset. The clear and consistent separation of expression patterns between the two cell types confirms that the selected "Gold Standard" genes represent cell type–specific markers and validates their use as a reference set for benchmarking integration performance.
