## SupplementaryTablesS1-S5 for "Characterizing and Mitigating Protocol-Dependent Gene Expression Bias in 3′ and 5′ Single-Cell RNA Sequencing"

**Table S1.** The original cell-type annotations for Andrews dataset and their respective coarse cell types used for further analysis. If the cell-type name is not in the table, its original representation was preserved.

| **Original Cell-Type Annotations** | **New Cell-Type Annotations** |
|---|---|
| CD4-positive, alpha-beta T cell,<br>CD8-positive, alpha-beta T cell,<br>natural killer cell | T/NK cell |
| centrilobular region hepatocyte,<br>midzonal region hepatocyte,<br>periportal region hepatocyte,<br>hepatocyte | hepatocyte |
| classical monocyte,<br>Kupffer cell,<br>monocyte,<br>macrophage | monocyte/macrophage |
| endothelial cell of artery,<br>endothelial cell of hepatic sinusoid,<br>endothelial cell of pericentral hepatic sinusoid,<br>endothelial cell of periportal hepatic sinusoid,<br>vein endothelial cell | endothelial |

**Table S2.** The original cell-type annotations for Suo dataset (except Suo thymus) and their respective coarse cell types used for further analysis. If the cell-type name is not in the table, its original representation was preserved.

| **Original Cell-Type Annotations** | **New Cell-Type Annotations** |
|---|---|
| B-1 B cell,<br>B-2 B cell,<br>mature B cell,<br>immature B cell | B cell |

| | |
|---|---|
| CD8-alpha-alpha-positive,<br>alpha-beta intraepithelial T cell,<br>double-positive,<br>naive thymus-derived CD4-positive,<br>alpha-beta T cell,<br>naive thymus-derived CD8-positive,<br>regulatory T cell,<br>T cell<br>double negative thymocyte,<br>alpha-beta thymocyte | T cell |
| common dendritic progenitor,<br>dendritic cell,<br>plasmacytoid dendritic cell,<br>pre-conventional dendritic cell | dendritic cell |
| common myeloid progenitor,<br>early lymphoid progenitor,<br>granulocyte monocyte progenitor cell,<br>hematopoietic multipotent progenitor cell,<br>hematopoietic stem cell,<br>megakaryocyte-erythroid progenitor cell | hematopoietic progenitor cell |
| fraction A pre-pro B cell,<br>large pre-B-II cell,<br>late pro-B cell,<br>pro-B cell,<br>small pre-B-II cell | B cell progenitors |
| granulocyte,<br>myelocyte,<br>neutrophil,<br>promyelocyte | granulocyte |
| group 3 innate lymphoid cell,<br>group 2 innate lymphoid cell,<br>innate lymphoid cell | innate lymphoid cell |
| Kupffer cell,<br>Macrophage,<br>monocyte,<br>promonocyte | monocyte/macrophage |
| smooth muscle cell,<br>vascular associated smooth muscle cell | smooth muscle cell |

**Table S3.** The original cell-type annotations for Leader dataset and their respective coarse cell types used for further analysis. If the cell-type name is not in the table, its original representation was preserved.

| **Original Cell-Type Annotations** | **New Cell-Type Annotations** |
|---|---|
| dendritic cell,<br>conventional dendritic cell,<br>CD1c-positive myeloid dendritic cell | dendritic cell |
| epithelial cell of lung,<br>club cell,<br>multiciliated epithelial cell,<br>pulmonary alveolar type 1 cell,<br>pulmonary alveolar type 2 cell | respiratory epithelial cell |
| bronchus fibroblast of lung,<br>fibroblast of lung | fibroblasts |
| smooth muscle cell,<br>pericyte | smooth muscle/pericyte cell |
| endothelial cell of lymphatic vessel,<br>capillary endothelial cell,<br>pulmonary artery endothelial cell,<br>vein endothelial cell | endothelial cell |

**Table S4.** The original cell-type annotations for Jardine dataset and their respective coarse cell types used for further analysis. If the cell-type name is not in the table, its original representation was preserved.

| **Original Cell-Type Annotations** | **New Cell-Type Annotations** |
|---|---|
| precursor B cell,<br>immature B cell,<br>naive B cell | B cell |
| fraction A pre-pro B cell,<br>pro-B cell | B cell progenitors |
| CD4-positive, alpha-beta T cell,<br>CD8-positive, alpha-beta T cell,<br>regulatory T cell,<br>mature NK T cell,<br>CD16-negative, CD56-bright natural killer cell,<br>human, | T/NK cell |

| | |
|---|---|
| natural killer cell,<br>immature natural killer cell | |
| common dendritic progenitor,<br>dendritic cell | dendritic cell |
| myelocyte,<br>neutrophil,<br>promyelocyte,<br>eosinophil,<br>basophil | granulocyte |
| endothelial cell of sinusoid,<br>endothelial tip cell,<br>endothelial cell | endothelial cell |
| muscle cell,<br>muscle precursor cell,<br>precursor cell,<br>chondrocyte,<br>fibroblast,<br>myofibroblast cell | stromal/mesenchymal lineage |
| primitive red blood cell,<br>erythrocyte | RBCs |

**Table S5.** Description of cell-type sparsification strategy used for the biological Use Case. Each column indicates which cell types were retained for each protocol.

| **Dataset** | **Only 3′** | **Only 5′** | **Both 3′ and 5′** |
|---|---|---|---|
| Park (thymus) | CD8-positive, alpha-beta T cell,<br>gamma-delta T cell,<br>T-helper 17 cell,<br>double negative thymocyte,<br>thymocyte | regulatory T cell,<br>T cell,<br>double-positive, alpha-beta thymocyte | CD4-positive, alpha-beta T cell,<br>alpha-beta T cell |
| Suo (liver) | dendritic cell,<br>endothelial cell,<br>granulocyte | natural killer cell | B cell progenitors,<br>B cell,<br>hematopoietic progenitor cell,<br>innate lymphoid cell, |

| | | | |
|---|---|---|---|
| | | | T cell |
| Jardine (bone marrow) | dendritic cell,<br>early lymphoid progenitor,<br>macrophage,<br>promonocyte,<br>RBCs | B cell,<br>mast cell,<br>megakaryocyte,<br>osteoclast | granulocyte monocyte progenitor cell,<br>B cell progenitors |
